# Characterization of hybrid hydrogels combining natural polymers

**DOI:** 10.1101/2023.09.07.556716

**Authors:** Derin Taylan, İlayda Güneş, Emine Durukan, Özgül Gök

**Affiliations:** Researcher at Acibadem University; Acibadem University

**Keywords:** Hydrogels, Gelatin, PEG

## Abstract

Hydrogels have emerged as an important field of study in biomedical engineering, biomaterials research and soft tissue modeling due to their versatility and customizable structures. They are three-dimensional networks capable of absorbing significant amounts of water and can be chemically or physically bonded, rendering them insoluble. The unique properties of hydrogels, such as excellent biocompatibility, degradation abilities, and ability to form chemical bonds between macromolecules, make them valuable in a variety of applications. One of the main applications of hydrogels is tissue engineering, where they are used in implantation, surgical procedures, targeted drug delivery and tissue regeneration. Hydrogels can act as scaffolds to mimic the properties of natural tissues, including mechanical strength, degradation properties, gel transformation, and swelling. Crosslinking hydrogels with other polymers, both natural and synthetic, allows their degradation rate to be controlled. Composite hydrogels or biohybrid gels offer advantages such as consistent properties between batches and a high degree of control in the manufacturing process. Using different types of polymers, researchers can fine-tune the chemical and physical properties of the resulting hydrogel, which is crucial for certain biomedical applications such as drug delivery, biocompatibility, and mechanical stability. In this study, our aim is to make hybrid hydrogel scaffolds more suitable for biomedical applications, which can show the healing effect of adding natural polymers to the scaffold structure. Our results revealed that the hybrid hydrogel obtained by chemically linking gelatin polymer chains to PEG polymers during crosslinking via UV light provides more advantageous properties in the final hydrogel structure, such as better durability and pore sizes for future cell placement in tissue engineering applications.

## 1. INTRODUCTION

Hydrogels have emerged as a fascinating and versatile class of biomaterials over the last few decades. They were first described in the 1960s, and since then, researchers have been exploring their potential applications in various fields, particularly in biomedicine. Hydrogels are three-dimensional polymeric structures with unique properties, such as swelling and collapse capabilities, flexibility, biodegradability, biocompatibility, and softness. [1]

Hydrogels have been at the forefront of biomedical engineering, biomaterials research, and soft tissue modeling due to their great manipulation/ tailoring potential. They can be defined as high water absorbing, chemically or physically bonded insoluble three-dimensional (3D) networks. They offer great benefits such as excellent biocompatibility, degradation, high water absorption, and forming chemical linkages between macromolecules. These macromolecules forming hydrogels could later be used to create scaffolds, mimicking tissue when they are found in a stable form. Additionally, scaffolds may acquire different characteristics: mechanical strength, degradation properties, gel conversion, and swelling when crosslinked with other polymers. [1,2]

In tissue engineering, hydrogels are used in applications involving implantation, surgical and other invasive procedures to remove otherwise undegradable materials, targeted drug delivery, or cells and tissue regeneration [2].It is important to consider the effect of porosity and pore size in fine-tuning the degradation rate of hydrogels for specific applications. Although varying parameters such as the type and molecular weight of polymers or the degree of crosslinking can allow regulation of degradation, in vivo cellular and enzymatic processes can significantly affect these degradation rates.” Thus, it acts as a minimizing permeation and affects degradation capabilities, rendering it one of the more significant properties of hydrogels. [3]

Natural and synthetic polymers are crosslinked to form composite hydrogels to control the degradation rates of hydrogels. In addition, composite hydrogels, or biohybrid gels, offer advantages such as improved batch-to-batch consistency and a high degree of control during the fabrication processes. In short, using differ-ent types of polymeric materials allows for fine-tuning of chemical and physical properties within the resultant hydrogel, which may be critical to drug partitioning, physiological compatibility, chemical stability, environmental response, and mechanical integrity.[4,5,6]

The ability of hydrogels to absorb water is attributed to hydrophilic functional groups attached to the polymeric backbone, while their resistance to dissolution is due to cross-links between network chains. The structure and properties of hydrogels can be modified by varying the concentrations, structures, functionalities of the monomers, and the cross-linkers used during synthesis. There are two main types of hydrogels: physically cross-linked and chemically cross-linked. Physically cross-linked hydrogels form through non-covalent interactions, while chemically cross-linked hydrogels are held together by covalent bonds, which provide higher stability.Efforts have been made to synthesize hydrogels with improved mechanical properties and “intelligence,” where they can respond to external stimuli like pH, temperature, electricity, light, or biological molecules like enzymes. This responsiveness expands their potential applications, such as in controlled drug delivery systems, biosensors, tissue engineering scaffolds, and actuator devices.[6]

In this study, our aim was to prepare hybrid hydrogel scaffolds which could show the ameliorating effect of addition of natural polymers into scaffold structure so that it becomes more suitable for biomedical applications. Herein, a hydrogel only made out of a very hydrophilic yet synthetic polymer PEG (Polyethylene glycol) was first synthesized by UV-curing ability of methacrylate groups in the polymer structure. The PEG chains were crosslinked into each other with the reaction of methacrylate groups via a radical-based polymerization strategy. The obtained hydrogel was improved by its both mechanical and morphological properties by the addition of a naturally occurring biopolymer, gelatin, into its structure. By this way, hybrid hydrogels were prepared and demonstrated to provide better characteristics for further biomedical applications. Moreover, the incorporation of gelatin polymer into PEG-based hydrogels was studied both ways: physical entrapment of gelatin chains into pores of the hydrogel by simple blending and chemical immobilization of gelatin chains from its functional methacrylate units into obtained hydrogel structure with the help of UV curing ability of methacrylate groups. Our results revealed that the hybrid hydrogel obtained by chemical bonding of gelatin polymer chains to the PEG polymers during the crosslinking via UV light provided more advantageous properties in the final hydrogel structure, like better durability and pore sizes for future cell accommodation in tissue engineering applications.

## 2. MATERIALS AND METHODS

### 2.1. Materials

Gelatin (Type B, powder, BioReagent, suitable for cell culture), Methacrylic Anhydride (contains 2.000 ppm topanol A as inhibitor, 94%) were purchased from Sigma Aldrich for the synthesis methacrylate anhydride conjugated polymers.

Poly(ethylene glycol) dimethacrylate (average Mn 750, contains 900-1100 ppm MEHQ as inhibitor), Irgacure 2959 (2-Hydroxy-4^′^-(2-hydroxyethoxy)-2-methylpropiophenone) were purchased from Sigma Aldrich for preparation of crosslinked polymer.

Carbonate Buffer was prepared from calcium carbonate and calcium bicarbonate (Sigma Aldrich) in distilled water to dissolve gelatin polymer.

## 2.2. Methods

### 2.2.1. Preparation of UV-curable PEGdiMA Hydrogel

Briefly 500 μL PEGdiMA and 5 mg Irgacure was mixed by continuously stirring at 50 °C and 1500 rpm until Irgacure being dissolved homogeneously. After being dissolved, it was put under the UV light (365 nm) for 5 minutes until being crosslinked. It was washed with distilled water to remove side products. After that hydrogel was stored at −80 °C for 1 h and lyophilized to obtain dry PEGdiMA polymer.

### 2.2.2. Preparation of UV-curable Gelatin-PEGdiMA Hydrogel

Briefly 10 mg Gelatin was dissolved in 375 μL Carbonate Buffer (CB) and then 125 μL PEGdiMA was added. And then 5 mg Irgacure was added and continuously stirred at 50 °C and 1500 rpm until Irgacure was dissolved homogeneously. After being dissolved, it was put under the UV light (365 nm) for 5 minutes until being crosslinked. It was washed with distilled water to remove side products. After that, hydrogel was stored at −80 °C for 1 h and lyophilized to obtain dry Gelatin-PEGdiMa polymer.

### 2.2.3. Preparation of UV-curable GelMA-PEGdiMA Hydrogel

Briefly 5 mg Gelatin-Methacrylate (GelMA) was dissolved in 375 μL distilled water and then 125 μL PEGdiMA was added. And then 5 mg Irgacure was added and continuously stirred at 50 °C and 1500 rpm until Irgacure was dissolved homogeneously. After being dissolved, it was put under the UV light (365 nm) for 5 minutes until being crosslinked. It was washed with distilled water to remove side products. After that, hydrogel was stored at - 80 °C for 1 h and lyophilized to obtain dry GelMA-PEGdiMA polymer.

## 3. RESULTS AND DISCUSSION

### 3.1. Swelling

Swelling is a characterization method used to determine the gel conversion rates of the polymers when interacted with water. It enables prediction on the release rates of the active ingredients which may be placed inside the scaffold for drug delivery and is controlled by stress relaxation in the swelling polymer network. Swelling results are often used to interpret the nutrient and metabolite transport efficiency of the scaffold without negatively impacting the mechanical stability. In short, hydrogels should be porous enough to be biocompatible while also being efficient upon degradation and delivering the required substance. Material characterization is reached through the polymer’s swelling capacity and absorbent properties, which is important for predicting their performance and behavior during application. Consequently, applications such as drug release depend highly on swelling properties due to their capability to ensure that the drug molecules penetrate further into the polymer matrix and that they have a controlled release. In our study, we observed the swelling results of H2 and H3 to see the effects of synthetic polymers and natural polymers. The data showed that H3 had better swelling capacity than H2. (H3 was not homogenois so it degraded)

**Tablo 1.** Swelling Data. (**:degraded)

| Exp<br>T | H2 (mg) | H3 (mg) |
| --- | --- | --- |
| 0 | 105,0 | 96,0 |
| 1 | 155,47 | 226,9 |
| 2 | 172,2 | 302,7 |
| 5 | 213,2 | 335,8 |
| 10 | 218,9 ** | 397,8 |
| 15 | 234,4 | 425,7 |
| 30 | 262,2 | 366,8 |
| 45 | 281,5 | ** |

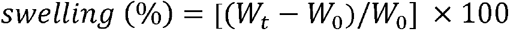

***Formulation 1. Swelling Formulation*.*(Wt is the weight of the gel for a given time. Wo is the initial weight of the gel*.)**

Example calculation: First, we need to find the initial weight (Wo) of the gel. When we see the value 0 in the “Time/Exp no” column for the initial time point, we find the values 96.0 and 105.0 in the Exp 1 and H2 columns, respectively. Suppose this corresponds to the weight of the beginning.

Next, we find the weight (Wt) of the gel for the other time points and calculate the gel swelling. For example, for time point 1 we see 226.9 in the Exp 1 column. In this situation:

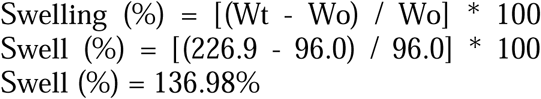

**Tablo 2.** Swelling ratio.

| Exp<br>T | H2 | H3 |
| --- | --- | --- |
| 0 | W0=105,0 | W0=96,0 |
| 1 | 47,43% | 136,46% |
| 2 | 64,00% | 215,31% |
| 5 | 102,09% | 249,58% |
| 10 | 129,58% | 313,75% |
| **15** | 145,71% | 344,06% |
| **30** | 149,14% | 281,67% |
| **45** | 168,09% | - |

**Graph 1.**
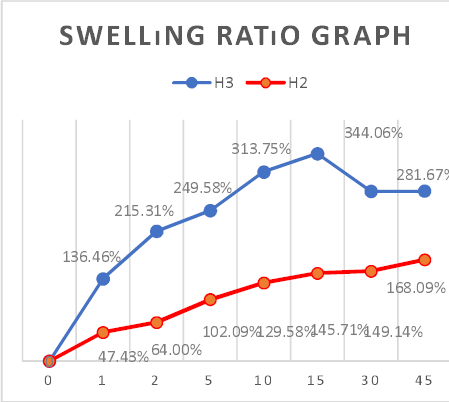
Swelling Ratio graph.

Hydrogel made up of PEGdiMA and gelatin had less swelling capacity indicating less mechanical stability with 149,14% at 30 min.

### 3.2. Rheometer

Rheometer is a device that is used to obtain deformation, flow, viscosity, etc. properties of materials to understand better the coupling between the shear flow and the material microstructure and its organization at the supramolecular scale. Changes in the microstructure produce changes in the deformation rates of polymers, such as during shear thinning when a great deal of reduction is observed in the viscosity rates of the polymers. In this study, an oscillation test was made to examine the deformation rates of hydrogels.

The degradation behavior of obtained hydrogels was examined using rheometry. Here, G^″^ represents viscosity (loss modulus), and G^′^ represents elasticity (storage modulus). Initial states display the elastic properties of samples. After a while, G^″^ and G’ overlap and split. The overlapping point is where the deformation in the hydrogel structure happens and where the degradation starts. After this point, while the value of G’ decreases, the value of G’’ increases relatively more than G’, which means that the liquid-like properties of hydrogels increase.

When these two rheometer results are compared, the value level of G’’ belonging to ‘H2’ is lower than G’’ of ‘H3’ as seen. This difference means the liquid-like property is much more in ‘H3’ than ‘H2’, and the swelling ratio is better at ‘H3’, which supports the data collected for swelling ratios. Thus, the overlapping point is almost the same for the two samples, which means that the degradation rate of these two samples happens at the same frequency.

**Figure 1:**
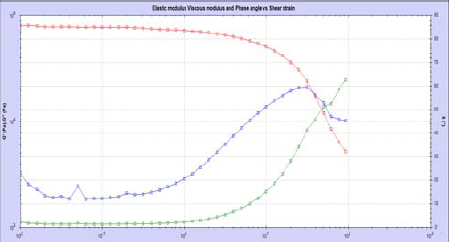
Rheometer test for H2.

**Figure 2:**
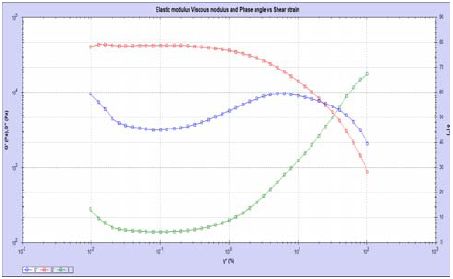
Rheometer test for H3.

### 3.3. FT-IR

In order to make a functional group analysis, 500 μL of the PEGdiMA, Gelatin-PEGdiMA and GelMA-PEGdiMA were lyophilized. A certain amount of dried samples were used for FT-IR Spectroscopy analysis. The ability to associate certain absorption bands, also known as group frequencies, with particular molecular components makes it easier to understand spectra. Depending on the type of group frequency, the mid-infrared region is separated into 4 areas. The peak seen between 2000 cm^-1^ and 1500 cm^-1^ is quantified as the range of double bonds. The main bands in this range are stretching vibrations of C=O bonds. The stretching vibrations of the OH and NH are mostly produced in the ranges between 3400 and 3300 cm^-1^ and 3650 to 3250 cm^-1,^ respectively. The analysis verified the information about the molecules present in gelatin.

**Figure 3.**
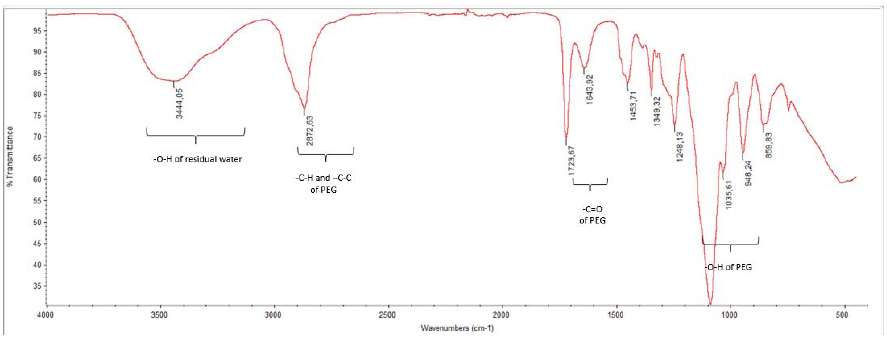
FT-IR spectra of H1.

**Figure 4.**
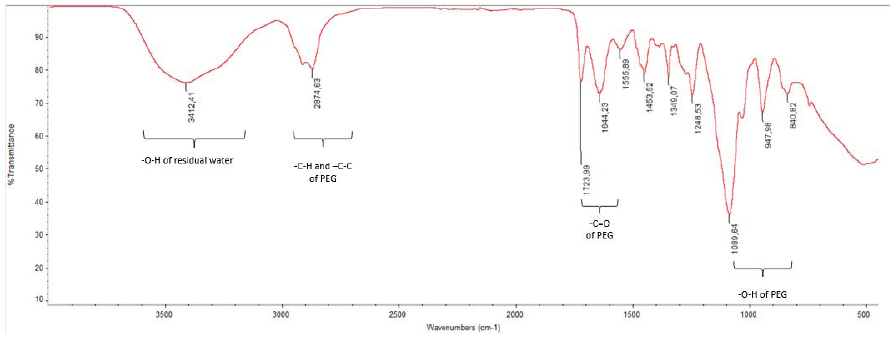
FT-IR spectra of H2.

**Figure 5.**
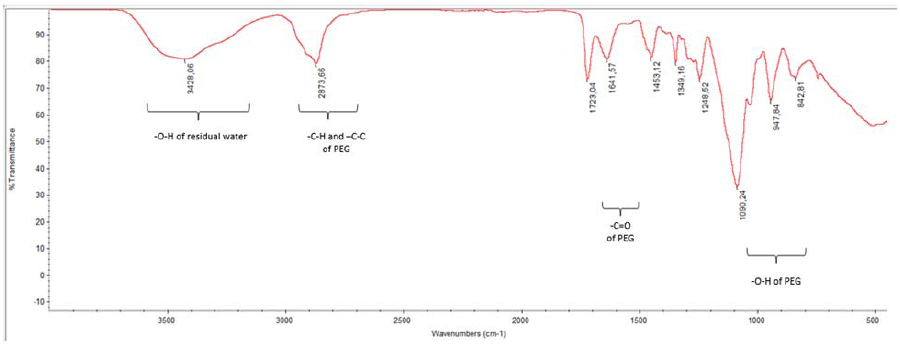
FT-IR spectra of H3.

### 3.4. SEM

A scanning electron microscope (SEM) is an electron microscope that produces images of a sample by scanning the surface with a focused beam of electrons. The electrons interact with atoms in the sample, producing various signals containing information about the sample’s surface topography and composition.

In this study, looking at the SEM images, porous structures were observed in the hydrogels, which would be expected of them to provide cell migration, proliferation, and emplacement. Images of H2 and H3 indicate that they are about 2500 microns and could easily host cells (10-15 microns) for further biomedical applications. However, it is also evident that H3 has a more porous structure which would render it more effective as a scaffold.

**Figure 6:**
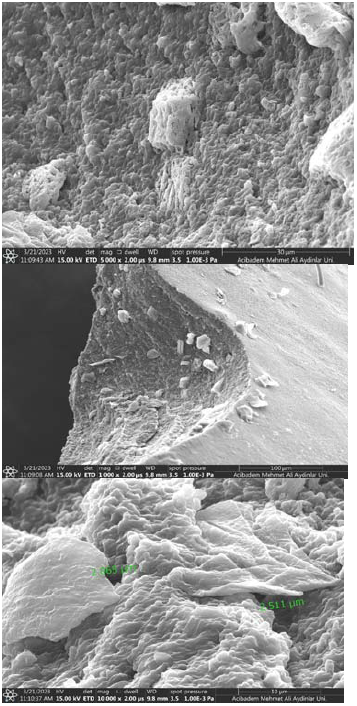
SEM images for H2 hydrogel.

**Figure 7:**
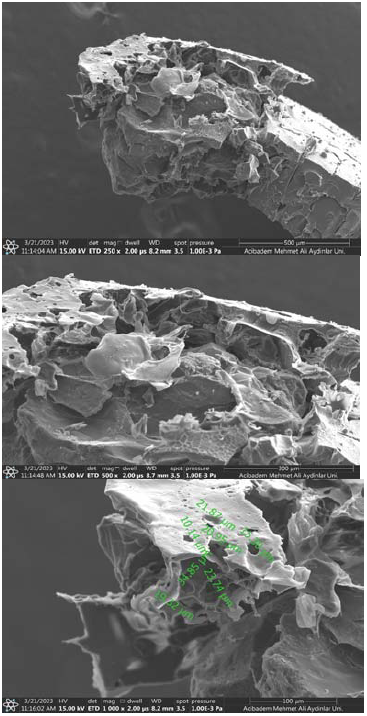
SEM images for H3 hydrogel.

## CONCLUSIONS

This study aimed to increase the biomedical application efficiency of hydrogels by combining natural polymers with synthetic polymers. They used four different methods to analyze and compare the properties of hydrogels: scanning electron microscopy (SEM), swelling behavior, rheometry and Fourier-transform infrared spectroscopy (FT-IR).

H2 was formed by blending natural gelatin polymers with synthetic PEG polymers.H3 was obtained by chemical bonding of gelatin polymer chains to PEG polymers during crosslinking using UV light. The analysis results showed that the hybrid hydrogel (H3) obtained by chemical bonding of gelatin to PEG exhibited more advantageous properties in the final hydrogel structure compared to pure PEG hydrogel (H1), such as improved durability and pore sizes suitable to accommodate cells in tissue engineering applications. When the swelling behavior was examined, H3 showed a 130% higher swelling ratio compared to H2, indicating a higher water absorption capacity. SEM images also supported this finding by showing that H3 was much more porous than H2. Rheometry data showed no significant difference in degradation rates between H2 and H3, suggesting that the chemical bonding of gelatin to PEG does not significantly affect the degradation behavior of the hydrogel.FT-IR analysis confirmed the presence of similar compounds in all three hydrogel samples, with the exception of the varying permeability of the C=O bond of PEG. This indicated that the chemical composition of the hydrogels was similar, but the presence of gelatin in H2 and H3 may have affected some of the bonds in PEG. Based on these findings, it was concluded that the composition of Gel-MA and PEG(H3) is the best candidate among the three hydrogels for biomedical applications due to its improved physical and chemical properties, high swelling capacity and pore structure suitable for tissue engineering purposes.

## Notes

### Competing Interest Statement

The authors have declared no competing interest.

### Summary of Updates

Addition of new authors.

